# A Sensitive Yellow Fever Virus Entry Reporter Identifies Valosin-Containing Protein (VCP/p97) as an Essential Host Factor for Flavivirus Uncoating

**DOI:** 10.1101/814764

**Authors:** Harish N. Ramanathan, Shuo Zhang, Florian Douam, Jinhong Chang, Priscilla L. Yang, Alexander Ploss, Brett D. Lindenbach

## Abstract

While the basic mechanisms of flavivirus entry and fusion are understood, little is known about the post-fusion events that precede RNA replication, such as nucleocapsid disassembly. We describe here a sensitive, conditionally replication-defective yellow fever virus (YFV) entry reporter, YFVΔSK/Nluc, to quantitively monitor the translation of incoming, virus particle-delivered genomes. We validated that YFVΔSK/Nluc gene expression can be neutralized by YFV-specific antisera and requires known flavivirus entry pathways, including clathrin- and dynamin-mediated endocytosis, endosomal acidification, YFV E glycoprotein-mediated fusion, and cellular LY6E expression; however, as expected, gene expression from the defective reporter virus was insensitive to a small molecule inhibitor of YFV RNA replication. YFVΔSK/Nluc gene expression was also shown to require cellular ubiquitylation, consistent with recent findings that dengue virus capsid protein must be ubiquitylated in order for nucleocapsid uncoating to occur, as well as valosin-containing protein (VCP)/p97, a cellular ATPase that unfolds and extracts ubiquitylated client proteins from large macromolecular complexes. RNA transfection and washout experiments showed that VCP/p97 functions at a post-fusion, pre-translation step in YFV entry. Together, these data support a critical role for VCP/p97 in the disassembly of incoming flavivirus nucleocapsids during a post-fusion step in virus entry.

**IMPORTANCE:** Flaviviruses are an important group of RNA viruses that cause significant human disease. The mechanisms by which flavivirus nucleocapsids are disassembled during virus entry remain unclear. Here we show that the yellow fever virus nucleocapsid disassembly requires the cellular protein-disaggregating enzyme valosin-containing protein, also known as p97.

## INTRODUCTION

Flaviviruses are a large, genetically-related group of positive-strand RNA viruses that are classified as a genus, *Flavivirus*, within the family *Flaviviridae*, including several medically important, arthropod-borne human pathogens such as dengue virus (DENV), Japanese encephalitis virus, West Nile virus (WNV), yellow fever virus (YFV), and Zika virus (ZIKV) (1). Infectious virions are small (∼50 nm diameter), enveloped particles that display 180 copies of the envelope (E) glycoprotein and a small transmembrane protein, M, on their surface, surrounding an inner nucleocapsid composed of the viral capsid protein and an unsegmented RNA genome of ∼11-kb (2, 3). While the surface of flavivirus particles are relatively well-defined, the nucleocapsid lacks apparent symmetry or higher-order structure (2, 4).

Flavivirus infection initiates through interaction of the viral E glycoprotein with one or more host cell attachment factors that serve to concentrate the virions on cell surface, as well as virus entry receptors that have only been partially identified (5). Virus internalization occurs through clathrin- and dynamin-dependent receptor-mediated endocytosis (6). As internalized virus particles pass through endosomes they encounter low pH, which triggers rearrangement of the E glycoprotein, leading to fusion and release of viral nucleocapsids into the cytoplasm of infected cells (Fig. 1A).

**Figure 1.**
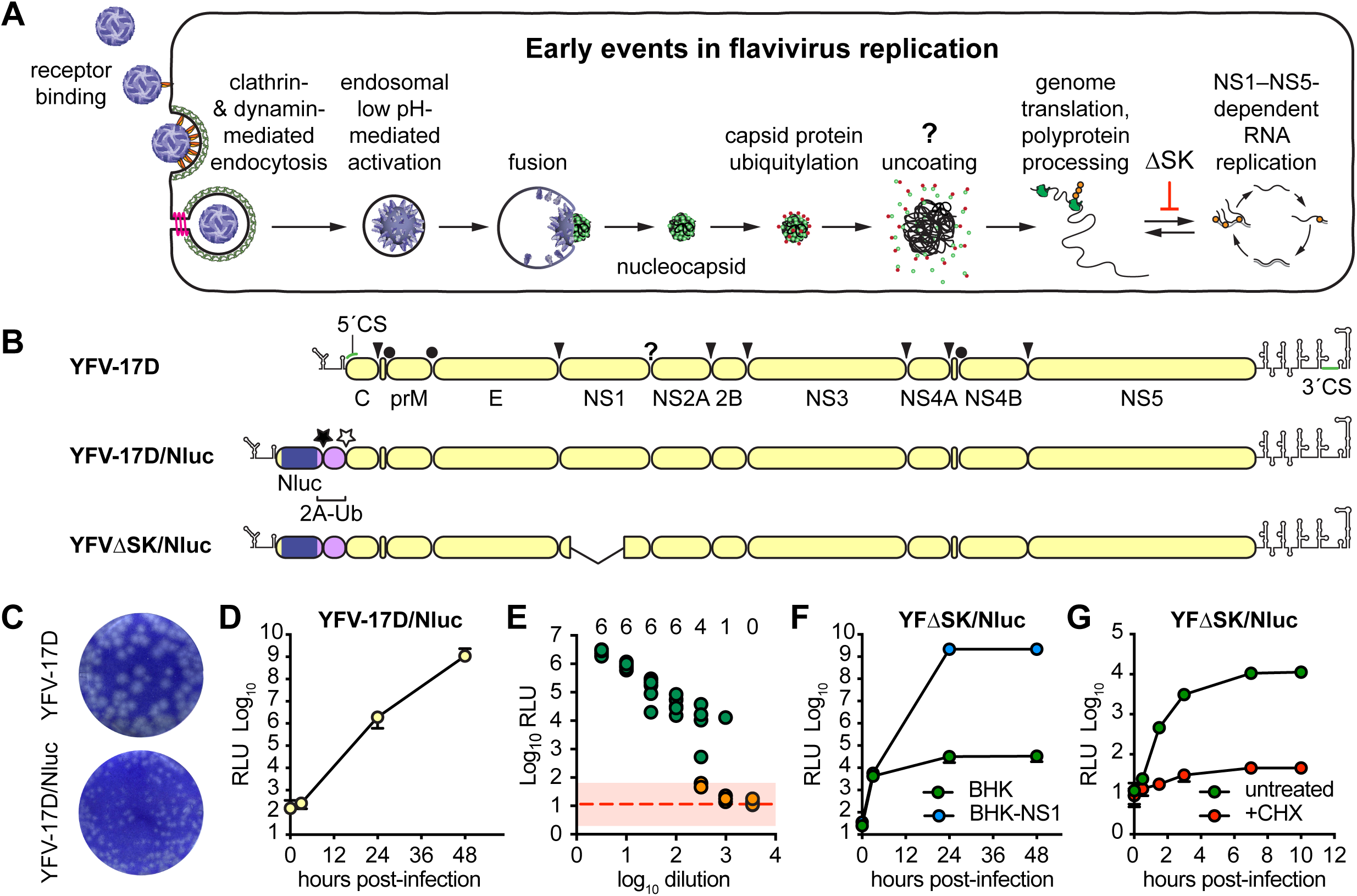
The Nluc reporter virus is a sensitive tool to monitor early events of flavivirus infection. (A) Early steps in the flavivirus life cycle, showing entry, uncoating, translation, and RNA replication. (B) Genome representation and polyprotein processing of YFV-17D, YFV-17D/Nluc and YFVΔSK/Nluc reporter viruses used in this study. Circles represent signal peptidase cleavage sites within the YFV polyprotein: downward arrowheads represent YFV NS2B-3 serine protease cleavage sites; the question mark represents the cleavage site of an unidentified cellular protease; filled and open stars represent the FMDV 2A translational skipping site and ubiquitin C-terminal hydrolase cleavage site, respectively. (C) Representative wells from YFV-17D and YFV-17D/Nluc plaque assays developed over 5 days with a 0.3% Avicel CL-611 overlay. (D) Time-course of Nluc expression after YFV-17D/Nluc infection (MOI 0.3) of BHK-21 cells. (E) Results of an endpoint dilution assay performed in six-fold replicate. Wells were scored as positive (green) or negative (orange) based on whether they were >2 σ (pink rectangle) away from the mean of uninfected controls (dotted red line); the numbers at the top of the graph represent the number of positive wells at each dilution. (F) Time-course of Nluc expression after YFV-ΔSK/Nluc infection (MOI 0.1) of BHK-21 or BHK-NS1 cells. (G) Time-course of Nluc expression after YFV-ΔSK/Nluc infection (MOI 0.1) of untreated or CHX-treated BHK-21 cells.

Once in the cytoplasm, flavivirus nucleocapsids are presumably disassembled to release the viral genome, which must be translated in order to initiate infection. The viral genome encodes a single open reading frame, which is translated to produce a polyprotein that is processed by viral and cellular proteases to yield three structural proteins: capsid (C), pre-M (prM), and E; as well as seven nonstructural (NS) proteins: NS1, NS2A, NS2B, NS3, NS4A, NS4B, and NS5 (Fig. 1B). NS1 is a secreted glycoprotein that is required for RNA replication; it forms dimers that remain peripherally associated with the inner leaflet of the endoplasmic reticulum (ER) membrane or on the cell surface, as well as secreted hexameric lipoprotein particles that induce strong humoral responses and contribute to flavivirus pathogenesis (3, 7). NS2A, NS2B, NS4A, and NS4B are polytopic membrane proteins required for RNA replication. NS2B and NS3 form a membrane-anchored enzyme complex with serine protease and RNA helicase activities essential for viral polyprotein processing and RNA replication (3, 8). NS5 is the viral RNA-dependent RNA polymerase and RNA capping enzyme (3, 9, 10). Once translated, the NS proteins presumably recruit the viral genome out of translation and into an RNA replication complex.

Little is known about the process of flavivirus nucleocapsid disassembly. Nucleocapsids obtained by solubilizing WNV particles with nonionic detergent are partially accessible for translation (11), suggesting that nucleocapsids may spontaneously uncoat. On the other hand, intact nucleocapsids can be isolated from detergent-solubilized tick-borne encephalitis virus particles (12); these nucleocapsids are stable in physiologic salt but dissociate in high salt, >0.5 M sodium chloride. In cell culture, however, DENV capsid protein must be ubiquitylated in order for nucleocapsid uncoating and genome translation to occur (13), suggesting that uncoating is an active process *in vivo*.

Here we describe a sensitive, conditionally replication-defective YFV reporter virus designed to probe the early, pre-replication events of the flavivirus life cycle. We validate the specificity of this reporter to monitor YFV entry and pre-replication events, confirm that YFV entry requires ubiquitylation, then used this tool to examine the hypothesis that YFV nucleocapsids are disassembled by valosin-containing protein (VCP), also known as p97.

VCP/p97 is a conserved and abundant eukaryotic AAA+ ATPase that uses the energy released by ATP hydrolysis to unfold ubiquitylated proteins and extract them from large macromolecular complexes (14, 15). VCP/p97 forms hexameric double-ring structures with a central pore (16, 17). Each subunit is composed of an N-terminal regulatory domain and two RecA-like ATPase domains that stack against each other to form the hexameric holoenzyme. VCP/p97 plays an essential role in protein homeostasis and genome stability; it is therefore an attractive target for anticancer therapies and several potent and specific VCP/p97 inhibitors have been developed (18). We show here that VCP/p97 inhibitors block the initiation of YFV infection, interfering with a post-fusion, pre-replication event during virus entry.

## RESULTS

### YFV reporter viruses are sensitive tools to monitor early events of flavivirus infection

A key challenge to studying the early life cycle events of some positive-strand RNA viruses is in detecting the initial rounds of translation of incoming, virion-released genomes. While viruses can be engineered to express sensitive reporter genes, signals produced by translation of the incoming viral genome are soon overwhelmed by translation of genomes produced by RNA replication. Therefore, in order to specifically study early, pre-replication events in the life cycle of flaviviruses, we sought to uncouple translation from RNA replication by constructing a YFV reporter, based on the live-attenuated strain 17D (YFV-17D), that is conditionally defective for RNA replication. The YFV-17D mutant YFVΔSK, which contains a large, in-frame deletion within the essential NS1 gene, is incapable of initiating RNA replication but can be complemented in *trans* (19). Thus, in the absence of NS1, a YFVΔSK-based reporter virus should allow viral entry, fusion, uncoating, and primary translation of the incoming genome to be monitored (Fig. 1A).

First, we constructed a full-length, infectious YFV-17D reporter virus that expresses the Nano-luciferase (Nluc) enzyme, based on previously described flavivirus reporter designs (20–22). We chose Nluc because of its smaller size (19.1 kDa; 171 codons), enhanced stability, and exquisite sensitivity compared to other luciferases (23). A cassette encoding Nluc, the foot-and-mouth disease virus 2A codon-skipping peptide, and a ubiquitin monomer was inserted in-frame into the YFV-17D infectious clone after the first 25 codons of the YFV-17D C gene, which contains the essential 5’ RNA cyclization sequence (24, 25), followed by the entire YFV polyprotein coding sequence, to generate YFV17D/Nluc (Fig. 1B). After transfection into BHK-21 cells, YFV-17D/Nluc RNA transcripts replicated and gave rise to infectious virus with peak titers similar to wild-type YFV-17D (≥1 × 10^7^ PFU/ml at 48 h post-transfection) but had a small plaque phenotype (Fig. 1C). Similar replication impairments have been reported with other flavivirus reporter constructs (22). Nluc expression was stably maintained for at least three serial virus passages in BHK-21 cells; we did not specifically address the long-term stability of the Nluc insert. Based on prior reports of flavivirus insert instability, we expect that Nluc expression will be lost with passage and therefore limited our experiments to early passage stocks of virus. Importantly, YFV-17D/Nluc was able to infect and replicate in BHK cells, as observed by the robust accumulation of Nluc activity over time (Fig. 1D). Robust Nluc expression was also observed upon YFV-17D/Nluc infection of other established cell lines, including Huh-7.5, HEK 293, SW-13, and primary mouse fibroblasts (data not shown). Furthermore, Nluc expression levels directly correlated with the amount of input virus in an endpoint dilution assay (Fig. 1E); notably, some replicates were Nluc-negative at higher dilutions, indicating that an end-point had been reached (i.e., some replicate wells received no virus, while other wells received one or a few viruses). Notably, Nluc activity was 10- to 100-fold higher in positive wells around the endpoint, which likely received only a single virus particle, than negative wells (Fig. 1E). Based on this, we were able to calculate a tissue culture-infectious dose, 50% endpoint (TCID_50_) of 3.54 × 10^4^ TCID_50_/mL, which was similar to the plaque infectivity titers (3.25 × 10^4^ PFU/mL) of this same early-harvest (18 h post-transfection), low-titer stock. We consistently noted that virus stocks harvested after cytopathic effects became evident contained significant Nluc activity, presumably due to enzyme release into the conditioned medium by cell lysis. Thus, early virus harvests provided optimal signal:noise ratio without requiring extensive washing during virus inoculation (see Methods for detailed information). Taken together, these data show that YFV-17D/Nluc is a sensitive reporter virus for measuring YFV-17D infection and replication.

Next, we generated a conditionally replication-defective construct, YFVΔSK/Nluc, containing a large in-frame deletion within the essential NS1 gene (19). Upon transfection of RNA transcripts into BHK-21 cells that express YFV NS1 (BHK-NS1), YFVΔSK/Nluc replicated, expressed Nluc, and produced infectious virus. YFVΔSK/Nluc virus infected BHK-NS1 cells and expressed robust Nluc activity (Fig. 1F, blue circles); however, infection of parental BHK-21 cells, which do not express NS1, led to modest but significant levels of Nluc expression (Fig. 1F, green circles). To confirm that the Nluc activity observed in BHK-21 cells that lack NS1 expression was due to the entry and translation of YFVΔSK/Nluc, we performed an additional time course in the presence or absence of cycloheximide (CHX), a potent inhibitor of translation. Nluc expression was detectable as early as 30 min post-infection, increased >10-fold higher than in CHX-treated cells by 90 min post-infection, and plateaued by 7 h post-infection (Fig. 1G). Given the tight requirement for NS1 in flavivirus RNA replication, these results demonstrate that YFVΔSK/Nluc is a sensitive reporter to measure the translation of incoming viral genomes.

### Nluc reporter viruses mimic authentic flavivirus infection and Nluc activity correlates with cellular entry

To further validate the utility of YFVΔSK/Nluc for monitoring early stages of virus infection, we asked whether it exhibited known features of flavivirus entry and replication. First, we asked whether Nluc expression was sensitive to YFV-specific neutralizing antibodies. As shown in Fig. 2A, Nluc expression from both YFV-17D/Nluc and YFVΔSK/Nluc was neutralized by serum from an interferon-α/β R^-/-^ mouse immunized with YFV-17D, while non-immune control serum did not neutralize Nluc expression. Similarly, serum from a human YFV-17D vaccinee neutralized both reporter viruses, while pooled human serum did not (Fig. 2B). Together, these data confirm that Nluc expression is dependent on infectivity of the YFV reporter viruses.

**Figure 2.**
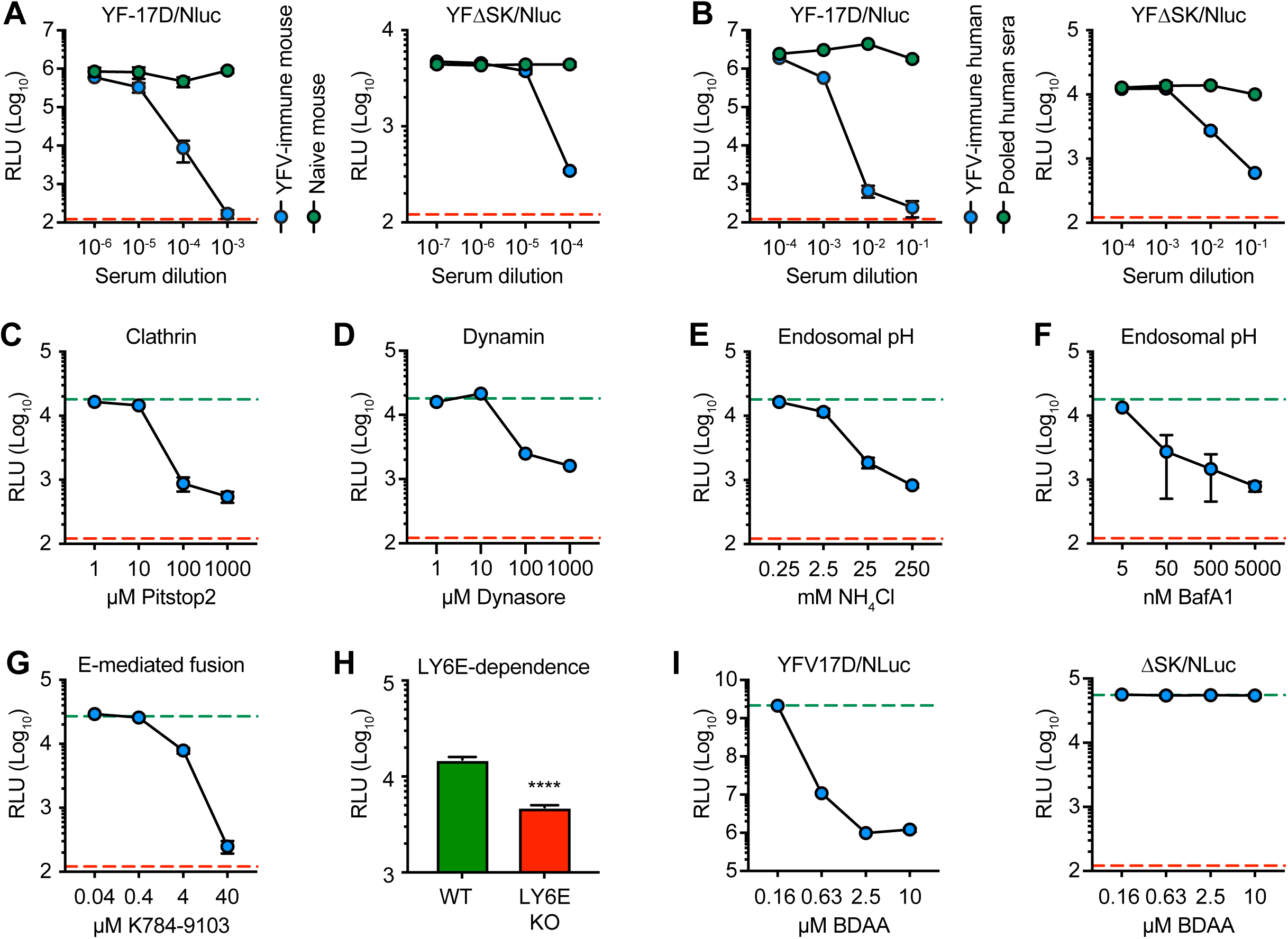
Nluc reporter viruses recapitulate authentic flavivirus infection. (A) Nluc expression after 24 infection with YFV-17D/Nluc (MOI 0.3; left panel) or YFVΔSK/Nluc (MOI 0.1; right panel) pretreated with YFV-immune or control mouse sera in BHK cells. (B) Nluc expression after 24 infection with YFV-17D/Nluc (MOI 0.3; left panel) or YFVΔSK/Nluc (MOI 0.1; right panel) pretreated with YFV-immune or pooled control human sera. (C) Nluc expression at 5 h post-infection with YFVΔSK/Nluc (MOI 0.1) in cells treated with Pistop2, a potent and specific inhibitor of clathrin-mediated endocytosis. (D) Nluc expression at 5 h post-infection with YFVΔSK/Nluc (MOI 0.1) of cells treated with Dynasore, a potent and specific inhibitor of dynamin. (E) Nluc expression from YFVΔSK/Nluc infection (MOI 0.1) of cells treated with NH_4_Cl, which buffers the endosomal compartment. (F) Nluc expression at 5 h post-infection with YFVΔSK/Nluc (MOI 0.1) of cells treated with Bafilomycin A, an inhibitor of endosomal acidification. (G) Nluc expression after 24 h infection with YFVΔSK/Nluc (MOI 0.1) treated with K784-9103, a potent and specific inhibitor of flavivirus fusion. (H) Nluc expression after 6 h infection of wild-type (WT) U2OS or LY6E knockout (KO) U2OS cells with YFVΔSK/Nluc (MOI 0.1). (I) Nluc expression after 24 h infection with YFV-17D/Nluc (MOI 0.3; left panel) or YFVΔSK/Nluc (MOI 0.1; right panel) in cells treated with K784-9103, a potent and specific inhibitor of YFV RNA replication. In all experiments, carrier control treatments are indicated by green dotted lines and the limit of Nluc detection (in parallel CHX-treated cells) by red dotted lines. All experiments were performed in triplicate except panel (H), which was performed in quadruplicate; error bars represent standard deviations from the mean.

Flaviviruses enter target cells via receptor-mediated endocytosis, which requires clathrin and dynamin, and delivery to endosomes, where viral fusion is induced by the low pH of this compartment (26). Consistent with this viral entry pathway, expression of Nluc by YFVΔSK/Nluc was sensitive to Pitstop2, an inhibitor of clathrin-coated pit formation (Fig. 2C), and to Dynasore, an inhibitor of dynamin (Fig. 2D). Furthermore, expression of Nluc by YFVΔSK/Nluc was sensitive to ammonium chloride (NH_4_Cl), a weak base that buffers endo-lysosomal compartments (Fig. 2E), and to Bafilomycin A1 (BafA1), an inhibitor of the vacuolar H^+^-ATPase pump (Fig. 2F). As shown in Fig. 2G, Nluc expression by YFVΔSK/Nluc was sensitive to K784-9103, a small-molecule that binds to DENV E protein and inhibits membrane fusion of DENV and other flaviviruses (27). Furthermore, Nluc expression was reduced 3.1-fold by genetic ablation of LY6E (Fig. 2H), a host factor that facilitates internalization of flaviviruses and other viruses (28, 29). These results confirm that YFVΔSK/Nluc gene expression are dependent on clathrin, dynamin, LY6E-mediated trafficking, endosomal acidification, and YFV E-mediated fusion, consistent with the known pathways of flavivirus entry.

We also examined the effects of benzodiazepine acetic acid (BDAA), a low micromolar inhibitor of YFV RNA replication that targets NS4B (30). In contrast to the entry-specific inhibitors used above, BDAA potently inhibited Nluc expression by YFV-17D/Nluc but had no effect on Nluc expression by YFΔSK/Nluc (Fig. 2I), confirming that YFΔSK/Nluc is a sensitive reporter of pre-replication events in the YFV life cycle.

### Ubiquitination and valosin-containing protein (VCP/p97) are essential for early stages of YFV infection

Since DENV genome uncoating requires a non-proteolytic ubiquitylation step (13), we next examined whether YFV also requires ubiquitylation at an early step in its life cycle. As shown in Figure 3A, expression of Nluc activity during YFVΔSK/Nluc infection was blocked by Pyr-41, a potent, specific, and irreversible inhibitor of the ubiquitin-activating enzyme E1 (31), confirming that ubiquitylation is required for an early, pre-replication event in the YFV life cycle. To clarify the stage in entry where ubiquitylation is required, we bypassed the viral entry process by directly transfecting YFVΔSK/Nluc RNA into BHK cells. As shown in Fig. 3B, Pyr-41 did not inhibit Nluc expression after RNA transfection. These results show that ubiquitylation is required for an early step in YFV entry, upstream of genome release (i.e., uncoating), consistent with the finding that ubiquitylation is required for the disassembly of the DENV nucleocapsid (13).

**Figure 3.**
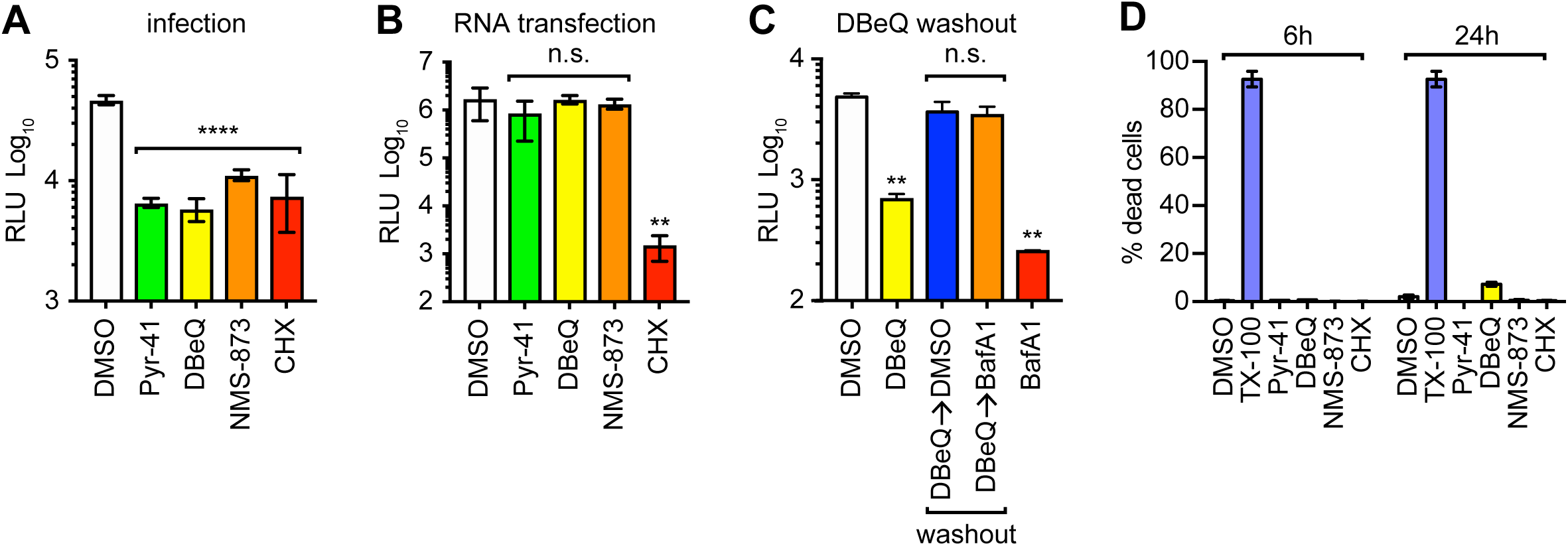
VCP/p97 is essential for an early, post-fusion stage of YFVΔSK/Nluc virus infection. (A) Nluc expression at 5 h post-infection with YFVΔSK/Nluc (MOI 0.1) of cells treated with Pyr-41 (50 *μ*M), DBeQ (10 *μ*M), NMS 873 (300 nM), or CHX (100 *μ*g/ml). (B) Nluc expression at 5 h post-transfection with 20 ng YFVΔSK/Nluc RNA in cells treated with the indicated compounds, as above. (C) Nluc expression at 6 h post-infection with YFVΔSK/Nluc RNA (MOI 0.1) in cells continuously treated with DMSO carrier control, DBeQ, or BafA1, as well as cells treated with DBeQ and subjected to washout as detailed in Methods. (D) Drug toxicity was quantified by plotting % dead cells against various inhibitor or control treatments, as above, at the indicated time points. All data correspond to averages from three independent experiments; error bars represent SD from the mean. Statistical significance was calculated by using ordinary one-way ANOVA (****, P<0.0001; ***, P<0.001; **, P<0.01; *, P<0.05; ns, not significant).

The above results suggested that ubiquitin can tag incoming nucleocapsids for subsequent uncoating by an unidentified host factor. The eukaryotic AAA+ ATPase VCP/p97 utilizes energy released from ATP hydrolysis to unfold ubiquitylated client proteins and extract them from larger complexes (14). We therefore tested the hypothesis that VCP/p97 promotes disassembly of the YFV nucleocapsid. As VCP/p97 is an abundant cellular protein, efficient knockdown takes several days to achieve a loss-of-function phenotype, which can cause pleiotropic effects on cells (32). Therefore, in order to specifically examine the role of VCP/p97 in YFV entry we chose two small molecules, DBeQ and NMS873, which work through different mechanisms of action to potently and specifically inhibit VCP/p97 in a matter of minutes, rather than days (33, 34). DBeQ and NMS873 potently inhibited Nluc expression after infection with YFVΔSK/Nluc virus particles (Fig. 3A) but did not inhibit Nluc expression after transfection of YFVΔSK/Nluc RNA (Fig. 3B), indicating that VCP/p97 is necessary for YFV entry, prior to the delivery and translation of incoming YFV genomes.

To clarify the step at which VCP/p97 functions during YFV entry, we conducted a washout experiment. As shown in Fig. 3C, DBeQ inhibition of YFVΔSK/Nluc entry could be reversed by drug washout at 1 h post-infection. Moreover, DBeQ washout bypassed the sensitivity to BafA1, indicating that VCP/p97 functions at a post-fusion step of YFV entry. We were unable to perform the converse experiment, washout of BafA1 followed by DBeQ treatment, because BafA1 washout was inefficient, consistent with the low nanomolar dissociation constant of this compound (35). Importantly, Pyr-41, DBeQ, NMS873, and CHX treatments were not toxic under the concentrations and time scales used in our studies (Fig. 3D), indicating that their abilities to block viral gene expression were not simply due to cellular toxicity. Taken together, these data suggest that VCP/p97 has a post-fusion, pre-replication function early in the YFV life cycle.

### VCP/p97 is required for early events in the life cycle of replication-competent flaviviruses

To determine whether ubiquitylation and VCP/p97 are also required for post-replication viral gene expression, we examined Nluc expression by the replication-competent YFV-17D/Nluc. As shown in Fig. 4A-B, Pyr-41 and DBeQ potently inhibited Nluc expression by 18 h post-infection with YFV-17D/Nluc, but not after 18 h post-transfection of YFV-17D/Nluc RNA. Since robust expression of Nluc activity at late times of infection (>8 h) depends on YFV-17D/Nluc replication (compare Figs. 1D, 1F, and 1G), these data suggest that ubiquitylation and VCP/p97 are specifically required for early events in the YFV life cycle.

**Figure 4.**
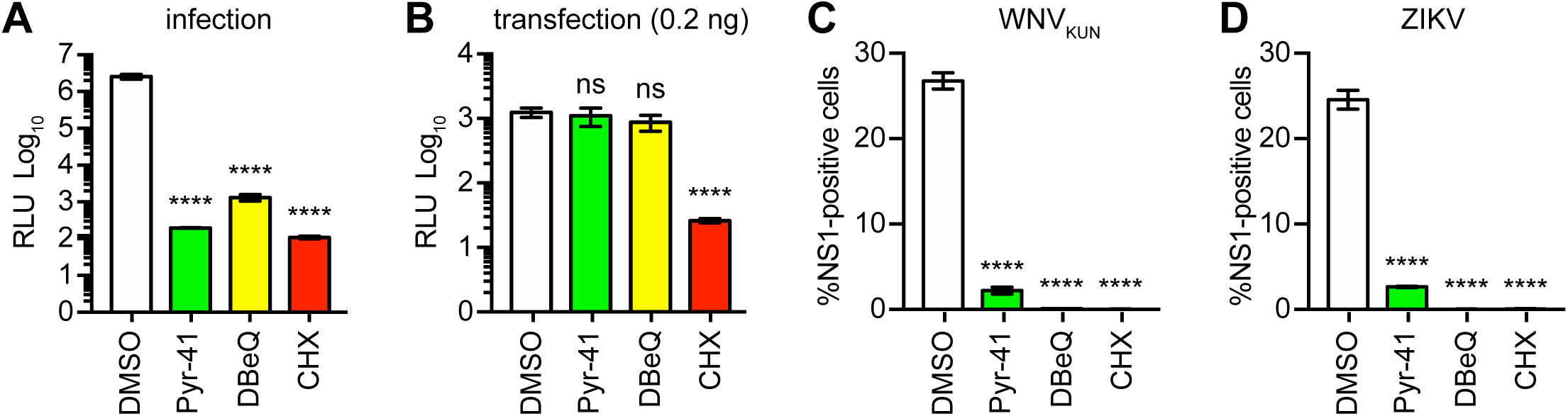
VCP/p97 activity is required for flavivirus infection. (A) Nluc expression at 5 h post-infection with YFV-17D/Nluc (MOI 0.3) of cells treated with the indicated compounds: Pyr-41 (50 *μ*M), DBeQ (10 *μ*M), NMS 873 (300 nM), or CHX (100 *μ*g/ml). (B) Nluc expression at 5 h post-transfection with 0.2 ng YFV-17D/Nluc RNA in cells treated with the indicated compounds, as above. (C) Infection of BHK cells with WNV_KUN_ (MOI 1.0) in cells treated with the indicated compound, as above. (D) Infection of BHK cells with ZIKV (MOI 1.0) in cells treated with the indicated compounds, as above. All data correspond to an average of three experiments; error bars represent standard deviation from the mean. Statistical significance was calculated by using one-way ANOVA (****, P<0.0001; ***, P<0.001; **, P<0.01; *, P<0.05; ns, not significant).

We next examined whether other flaviviruses depend on ubiquitylation and VCP/97 activity. As shown in Figs. 4C-D, Pyr-41, DBeQ, and CHX all potently inhibited the expression of NS1 at 24 h post-infection with the Kunjin strain of WNV (WNV_KUN_) or Cambodian FSS 13025 strain of ZIKV. These data suggest that multiple flaviviruses depend on cellular ubiquitylation and VCP/p97 activity.

## DISCUSSION

Flaviviruses initiate infection via a transient, multistep cellular entry process that poses some unique challenges to study. Early efforts focused on the identification of virus-specific receptors and in defining the mechanisms of antibody-mediated enhancement (36, 37). However, to date only a few flavivirus entry factors and putative entry receptors have been identified, including DC-SIGN (38, 39), DC-SIGNR (40), mannose receptor (41), and multiple phosphatidylserine receptors (42). Nevertheless, for many of these cell entry factors, the specific role(s) in flavivirus entry remains elusive (5).

The processes of flavivirus internalization and fusion were originally characterized by using biochemical and cell biological approaches with high MOIs of radiolabeled virus particles, revealing a requirement for endocytosis and endosomal acidification (43, 44). While working with infectious, radiolabeled virus particles may be slightly inconvenient, the use of high MOIs is potentially more problematic, as aggregates of virus particles can influence the apparent mechanisms of viral entry (45). More recently, the entry of individual, fluorescently-labeled flavivirus particles has been visualized at low MOI through live cell imaging (26, 46–48). An important caveat to this approach is that flavivirus preparations typically have relatively low specific infectivities (46, 47, 49), so it is difficult to know whether a given particle under observation is on a pathway toward productive infection.

Given the above considerations, we chose to pursue a function-based approach to study the productive entry of a recombinant YFV that expresses a reporter enzyme only after viral entry and translation. Several flavivirus reporter systems have been developed, mainly for high-throughput screening of viral replication (13, 22, 50–57). In one remarkable study, Byk, et al., adapted a *Renilla* luciferase-expressing DENV reporter virus to show that the incoming DENV capsid protein must be ubiquitylated prior to viral gene expression (13). However, an important consideration of this experimental design is that flavivirus-encoded reporter genes are continuously expressed, making it difficult to rigorously conclude that reporter activity was translated from an incoming viral genome versus viral RNAs produced by replication, i.e., because both viral entry and RNA replication can contribute to reporter gene expression, the translation of incoming genomes could only be inferred based on the kinetics of when reporter gene expression first became observable. Byk, et al. attempted to control for this concern by using CHX to inhibit translation (13); however, CHX inhibits viral gene expression irrespective of whether the genome was delivered by a virus particle or newly synthesized by RNA replication. A further consideration is that reporter enzymes differ in their specific activities; thus, enzymes with low specific activity require higher MOIs to achieve similar sensitivity of early translation events. In this regard, it is notable that Byk, et al. did not report the MOIs used in their DENV entry studies; however, several of their experiments used MOIs sufficiently high to allow incoming capsid protein to be detected by western blot (13). Despite these minor technical caveats, Byk, et al. clearly demonstrated that incoming DENV capsid protein must be ubiquitylated before viral gene expression can be detected.

Given the above considerations, we sought to build a sensitive YFV reporter specific for detecting translation of incoming viral genomes. First, we used the Nluc reporter gene, which exhibits >100-fold greater specific activity over firefly and *Renilla* luciferases (58). Second, we created a conditionally replication-defective reporter by incorporating a large, in-frame deletion in the essential NS1 gene, which can be supplied in *trans* (19, 59). The NS1 glycoprotein, which is expressed within the secretory pathway, contributes to the cytosolic process of RNA replication via interaction with the polytopic NS4A and NS4B membrane proteins, likely within the NS4A-2K-NS4B polyprotein intermediate (60–62). NS1 also has distinct membrane-alteration properties (61, 63), which likely contribute to replication complex assembly. In the absence of NS1 expression, flavivirus infections are halted prior to replication complex formation and the initial round of RNA synthesis (19, 61, 64, 65). Consistent with this, robust Nluc expression by YFV-17D/Nluc was sensitive to a YFV-specific RNA replication inhibitor, BDAA, while the modest Nluc expression by YFVΔSK/Nluc was not. Thus, the YFVΔSK/Nluc virus faithfully reports on early, post-fusion, pre-replication events in the flavivirus life cycle.

Our experimental approach should be generally applicable to other flaviviruses. It is notable that two groups previously described NS1 deletion-reporter virus systems for WNV and Omsk hemorrhagic fever virus (57, 66). These constructs were originally designed to reduce biosafety risks for high-throughput screening; our data suggest that these constructs should also be useful in dissecting early, pre-replication events in the life cycle of these flaviviruses. Further improvements to our design are also possible; for instance, smaller tags, such as a split Nluc reporter (55), could improve viral titers or allow post-fusion events to be monitored prior to viral genome translation.

We validated that YFVΔSK/Nluc gene expression was neutralized by YFV-specific antibodies and was dependent on several known pathways of flavivirus entry, including clathrin- and dynamin-mediated endocytosis (6, 26, 67–69), endosomal acidification (47, 68–71), E protein-dependent fusion (27), and dependence on LY6E (28, 29).

We then applied our YFV reporter system to address the role of ubiquitylation and protein homeostasis in flavivirus entry, which has been controversial. As part of a genome-wide RNAi screen, Krishnan, et al. first reported that knockdown of ubiquitin ligase CBLL1 inhibited internalization of WNV particles into HeLa cells, and that WNV entry was sensitive to proteasome inhibitors (72). However, these findings were called into question by Fernandez-Garcia, et al., who found that entry of WNV, YFV, and DENV was insensitive to rigorously validated knockdown of CBLL1 or by proteasome inhibitors (73). Furthermore, while JEV entry is also inhibited by proteasome inhibitors (74), these compounds can decrease the cellular pools of free ubiquitin (75–77), so the role of ubiquitylation vs. proteasome activity in flavivirus entry has been unclear. Byk, et al. brought clarity to this issue by demonstrating that ubiquitylation is required for DENV capsid disassembly (13). Furthermore, proteasome activity is dispensable for DENV entry, but is responsible for the turnover of incoming capsid protein, presumably after disassembly (13).

Our studies confirm that ubiquitylation is required for flavivirus entry, although the relevant substrate(s) are unknown. Given that incoming DENV capsid protein is turned over in a proteasome-dependent manner, and that inhibition of proteasome activity leads to the accumulation of a slightly higher molecular weight form of capsid protein (13), it is likely that YFV capsid is directly ubiquitylated. It is not yet clear how nucleocapsids are targeted for ubiquitylation, nor whether there is a preferred site on capsid protein for this modification. In this regard, DENV mutants lacking lysine residues in the capsid protein were able to infect and translate their genomes normally (13), suggesting that the capsid protein may be ubiquitylated at the N-terminus or other non-canonical residue(s) (78). Future work would be needed to identify of the relevant E3 ligase(s) and the type(s) of ubiquitin linkage that modify flavivirus capsid proteins.

VCP/p97 functions to unfold and extract proteins from macromolecular complexes in a ubiquitin- and ATP-dependent manner (14). For instance, VCP/p97 dissociates ubiquitylated IκBα from NF-κB, activating this transcription factor (79). VCP/p97 contributes to ER-associated degradation by extruding misfolded proteins from the secretory pathway for subsequent delivery to the proteasome (80). Similarly, VCP/p97 contributes to ribosome-associated quality control by extracting misfolded nascent polypeptides from the translation apparatus (81, 82). VCP/p97 has additional roles in extracting client substrates from chromatin, mitochondria, and other large macromolecular complexes. It is worth noting that VCP/p97 has a weak affinity for ubiquitin and relies on a large array of cofactors, which typically encode enzymatic activities to facilitate VCP/p97 substrate processing, or adaptor molecules, which simply link VCP/p97 to client substrates. Each of these adaptors and cofactors carry binding surfaces that recognize VCP/p97 and ubiquitin, respectively (83, 84). Thus, VCP/p97 contributes to diverse cellular functions based this modular cofactor- and adaptor-mediated targeting.

Based on our finding that VCP/p97 activity is required for an early, post-fusion event prior to the translation and replication of incoming YFV genomes, we propose a model wherein VCP/p97 functions to disassemble ubiquitylated nucleocapsids (Fig. 5). Although we have illustrated free nucleocapsids within the cytosol, we cannot exclude the possibility that fusion is tightly coupled to capsid ubiquitylation and disassembly, such that nucleocapsids may be disassembled as they are exposed to the cytosol. Consistent with our model, VCP/p97 activity was previously shown to be important for WNV, JEV, and DENV infections, although specific role(s) for VCP/p97 in virus entry were not determined (55, 85). We cannot also exclude a role for ribosome engagement to facilitate disassembly.

**Figure 5.**
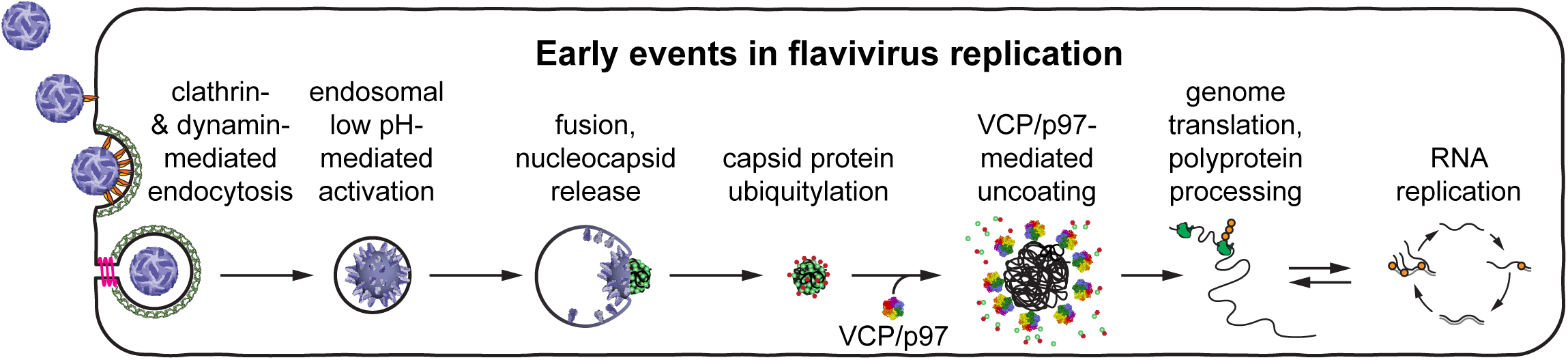
A revised model for the early, pre-replication events in flavivirus infection. Modified from Fig. 1A. Within the cytoplasm, capsid protein (green spheres) are ubiquitinated (small red spheres), leading to VCP/p97-dependent nucleocapsid disassembly and release of the viral genome prior to translation.

The nature of VCP/p97 interaction with flavivirus nucleocapsids is presently unknown but may well involve cellular VCP/p97 cofactors. It is possible that VCP/p97 may be directly recruited to endo-lysosomal organelles via binding to early endosomal autoantigen, clathrin, or caveolin, as demonstrated by previous proteomic studies (86–88). Once recruited to endosomes, VCP/p97 may have direct access to ubiquitylated capsid protein during fusion or to an unknown ubiquitylated host protein that associates with capsid protein.

## ACKNOWLEDGMENTS

We thank Dr. Charles M. Rice (The Rockefeller University) for providing pACNR/YF17D, pACNR/YFΔSK, pSINrep21/YF NS1, pCC1/YF17D, and pYF-17D/5ʹC25Venus2AUbi; Dr. John Schoggins (University of Texas Southwestern Medical Center) for providing LY6E knockout U2OS cells and parental U2OS cells; Dr. Marie Flamand (Institut Pasteur) for providing NS1-specific monoclonal antibodies; Dr. Sergei Kotenko (Rutgers University) for providing C57BL/6J IFN-α/β R-/- mice; and Dr. Doug Brackney (Connecticut Agricultural Experiment Station) for providing BHK-21 clone 15 cells. This work was supported by PHS grants AI087925, AI131518, and AI120113 (all to B.D.L), AI107301 (to. A.P.) and a Burroughs-Wellcome Fund Investigator in Pathogenesis Award (101539) to A.P.

## MATERIALS AND METHODS

### Cell lines and plasmids

Baby hamster kidney (BHK-21), clone 15 cells, BHK-21 cells stably expressing YFV-NS1 (BHK-NS1), WT U2OS cells, and LY6E-knockout U2OS cells were maintained in Dulbecco’s minimum essential medium (DMEM, Life Technologies, Inc., Gaithersburg, MD) supplemented with 10% heat-inactivated fetal bovine serum (FBS, Omega Scientific) and 1 mM nonessential amino acids. The construction and maintenance of pCC1/YF17D and pYF-17D/5ʹC25Venus2AUbi were previously described (20, 21). Plasmid pYF17D/Nluc was constructed by replacing the Venus coding region of pYF-17D/5ʹC25Venus2AUbi with that of the Nluc gene (Promega, Madison, WI) by using standard molecular biology techniques and verified by restriction digestion and sequencing. Briefly, the SrfI–NsiI region of pYF17D/5ʹC25Venus2AUbi was subcloned into a shuttle vector, pSL1180, generating the pSL1180/S17DN intermediate. The Nluc coding region was PCR amplified with primers YO-3008 (5’-ggg ccc GAG CTC ATG GTC TTC ACA CTC GAA GAT TTC GTT G-3’) and YO-3009 (5’-ggg ccc acc ggt CGC CAG AAT GCG TTC GCA CAG CCG CCA GC-3’) and Q5 polymerase (NEB), then cloned into the SacI and AgeI sites of pSL1180/S17DN, resulting in replacement of Venus with Nluc. The SrfI–NsiI fragment was then subcloned back into pCC1/YF-17D to generate pYF17D/Nluc. The construction and use of pYFΔSK as well as pSINrep21-NS1 was described previously (19, 64). To generate pYFΔSK/Nluc, the 7221-bp NsiI–ClaI fragment of pACNR/YFΔSK was subcloned into pYFV17D/Nluc cut with these same enzymes.

### Transfections and virus stocks

Small scale RNA transfections were performed by using *Trans*IT mRNA transfection reagent (Mirus Bio) according to manufacturer’s recommendations. BHK-NS1 stable cells were regenerated as previously described (19). Briefly, 1 µg pSINrep21/YFV NS1 plasmid DNA was transfected into BHK-21 clone 15 cells by using 8 µl *Trans*IT LT1 reagent (Mirus Bio, Wisconsin, MD) and low-serum OptiMem (Life Technologies). Transfected cells were then selected for one week in complete growth medium supplemented with 5 µg/ml puromycin. Reporter virus RNAs were transcribed from XhoI-linearized plasmid templates pYF17D/Nluc and pYFΔSK/Nluc by using SP6 RNA polymerase in the presence of the ARCA synthetic cap analog (New England Biolabs, Ipswich, MA). Primary stocks of reporter virus were generated by electroporation of BHK-21 clone 15 or BHK-NS1 cells with YFV RNA transcripts, as previously described (19). At 36 h post-electroporation, conditioned media containing primary YFV-17D/Nluc or YFVΔSK/Nluc stocks were harvested and clarified by centrifugation at 3,000 × *g* at 4°C for 10 min to remove cell debris.

Pilot experiments showed that primary stocks of YFV-17D/Nluc virus harvested at late time points (≥36 h post-transfection) contained high levels of Nluc released into the conditioned cell culture media, which correlated with the onset of virus-induced cytopathic effects. Therefore, to help minimize background, primary stocks of Nluc-expressing virus were dialyzed via ultrafiltration with Centricon plus-80 (containing 100-kDa nominal molecular weight cut-off polyethersulfone filters) to remove contaminating background Nluc activity. Furthermore, secondary virus stocks with minimal Nluc background were harvested at early (<24 h) times post-infection, albeit with reduced titers. Specifically, primary virus stocks were passaged on several 15-cm dishes of ∼70% confluent monolayers of BHK-21 or BHK-NS1 cells; after 1 h incubation at 37°C, the inoculum was removed and washed twice with complete DMEM, twice with PBS, then incubated in the presence of complete DMEM containing 2% FBS; conditioned cell culture media were harvested ∼18 h post-inoculation, clarified as before, and stored in 1 ml aliquots at −80°C. These early harvest virus stocks had very low contaminating Nluc activity that was effectively removed by washing infected cells three times with PBS.

### Virus infectivity

Infectivity measurements were determined by using plaque assays or endpoint dilution assays. Plaque assays were performed as previously described (89). For endpoint dilution assays with Nluc viruses, virus stocks were serially diluted in half-log (√10) intervals in DMEM containing 2% FBS and nonessential amino acids; each dilution was then added to BHK cells seeded in 96-well plates. At 24 h post-infection, cells were washed twice with complete DMEM, once with PBS, and Nluc activity was measured in all wells. Wells were scored positive if the Nluc activity was >2 σ from the mean of uninfected controls. Tissue culture infectious dose 50% endpoint (TCID_50_) values were calculated by using the method of Reed and Muench method, as previously described (60).

### Nluc activity

Nluc activity was measured by using the Nano-Glo luciferase assay (Promega). At indicated times in the assay, cells grown in 96-well plates were gently washed twice with complete DMEM and once with PBS, then lysed with 20 µl Nano-Glo luciferase assay buffer containing the substrate. Nluc activity was measured from cell lysates within 10 min of lysis (or substrate addition) by transferring lysate into white OptiPlate 96-well plates (PerkinElmer, Waltham, MA) and measured in a Berthold Centro XS3 LB 960 luminescent plate reader with readings integrated over 0.2 sec.

### Inhibitors and treatments

Ammonium chloride, Baf A1, cycloheximide, and DBeQ were purchased from Millipore-Sigma (Burlington, MA). Pitstop2 and Dynasore were purchased from Abcam, Cambridge, MA. Pyr-41 was purchased from MedChemExpress (Monmouth Junction, NJ). NMS 873 was purchased from Tocris (Minneapolis, MN). Except where noted, all drugs were added to cells 1 h prior to infection or RNA transfection at the following concentrations: Pyr-41 (50 *μ*M), DBeQ (10 *μ*M), NMS 873 (300 nM), and CHX (100 *μ*g/ml) and maintained in the cell culture medium throughout the experiments. The DENV fusion inhibitor K784-9103 was purchased from ChemDiv (San Diego, CA), and its structure and purity were confirmed by tandem liquid chromatography-mass spectrometry. Inhibition of E-mediated fusion was performed as previously described (27). Briefly, reporter viruses were incubated in medium containing the indicated concentrations of K784-9103, then mixed in a rotary shaker for 30 min at room temperature to allow inhibitor to bind to virus particles before adding to cells.

DBeQ washout experiments were performed by adding 0.1% DMSO, 10 µM DBeQ, or 50 nM BafA1 with YFVΔSK/Nluc virus (i.e., no pre-incubation). At 1 h post-infection, all cells were washed twice with complete medium, twice with PBS, and returned to media containing 0.1% (v/v) DMSO, 10 µM DBeQ, or 50 nM BafA1, as indicated; samples were collected 6 h later.

### Virus neutralization assay

C57BL/6J IFN-α/β R^-/-^ mice (90) were kindly provided by Dr. Sergei Kotenko, Rutgers University. Mice were bred in the Laboratory Animal Resource Center of Princeton University. All animal experiments were performed in accordance to protocol number 1930, which was reviewed and approved by the Institutional Animal Care and Use Committee (IACUC) of Princeton University. One female, six-month old mouse was infected intravenously with 1 × 10^7^ PFU of YFV-17D. At eighteen days post-infection, the mouse was boosted intravenously with 1 × 10^7^ PFU of YFV-17D; serum was collected seven days following this booster injection. Serum from a female non-infected C57BL/6J IFN-α/β R^-/-^ littermate was collected in parallel for use as a negative control. The YFV-immune human serum was obtained from a de-identified donor through the American Red Cross. Pooled human sera was purchased from Thermo-Fisher (Waltham, MA) as a negative control. For virus neutralization experiments, mouse and human sera were diluted into reporter virus stocks, and samples were incubated for 30 min at room temperature with tumbling. At the end of the incubation, samples were centrifuged briefly and 50 µl samples were added to 3 wells of BHK cells grown in 96-well plates; Nluc expression was measured after 5 h of infection.

### Cytotoxicity assay

Cells were seeded in 96-well plates and incubated with inhibitors at their respective experimental concentrations (indicated above) along with the membrane-impermeable nuclear dye Cytotox Green (Essen BioScience Inc. Ann Arbor, MI) and cell-permeable Hoechst stain (Sigma). Cytotoxicity was measured on an ImageXpress Pico (Molecular Devices) by quantifying the fraction of Cytox Green-positive, permeable cells among total cells.

### Immunostaining and FACS analysis

WNV_KUN_ and ZIKV NS1 were detected by immunostaining with 6B8-2D8, a flavivirus NS1 cross-reactive monoclonal antibody, originally raised against DENV-4 NS1 (a kind gift of Dr. Marie Flamand, Institut Pasteur). Briefly, cells were washed twice with PBS and treated with Accumax (Innovative Cell Technologies, Inc., San Diego, CA) to gently dissociate cells for FACS analysis. Dissociated cells were directly fixed in paraformaldehyde solution (2% (w/v) final) for 30 min at room temperature. Fixed cells were permeabilized with 0.2% saponin for ∼30 min on a rotating chamber, followed by two washes with PBS. Cells were incubated overnight with a 1:3000 dilution of antibody 6B8-2D8 in PBS containing 2% FBS, washed with PBS, and incubated with 1:1500 dilution of an anti-mouse secondary antibody conjugated to Alexa-680 fluorescent dye. Specificity of labeling was confirmed by parallel incubation with the dye-conjugated secondary Ab in the absence of primary antibody, and by performing the complete staining procedure on uninfected cells. At the end of incubation, cells were washed twice, resuspended in PBS and subject to FACS analysis to quantify the percent of NS1-positive cells by using the far-red channel.

